# Prenatal Environmental Determinants of Aromatase Brain-Promoter Methylation in Cord Blood: Chemical, Airborne, Pharmacological, and Nutritional Factors

**DOI:** 10.1101/2025.07.24.666701

**Authors:** Samuel Tanner, Katherine Drummond, Sarah Thomson, Kristina Vacy, Christos Symeonides, Boris Novakovic, Toby Mansell, Martin O’Hely, Richard Saffery, Mimi LK Tang, Peter D Sly, Peter Vuillermin, the BIS Investigator Group, Wah Chin Boon, Chol-Hee Jung, Daniel Park, Anne-Louise Ponsonby

## Abstract

Aromatase, an enzyme encoded by the gene *CYP19A1*, plays central roles in neurodevelopment. In the brain, its function is to convert androgens into neuroestrogens, ensuring balanced hormonal signalling. Both animal experiments and human studies have shown that, in males, disruption of aromatase, either genetically or epigenetically, can increase symptoms of autism. Prenatal exposure to bisphenol A (BPA), a common plastic chemical, can increase levels of DNA methylation—a key epigenetic modification—at the brain-specific *CYP19A1* promoter, P1.f, reducing *CYP19A1* expression. However, the extent to which other neurodevelopmentally relevant environmental exposures influence P1.f methylation remains unclear. Here, in the Barwon Infant Study (BIS) birth cohort (*N* = 906), we analysed the association between 25 prenatal exposures (from five classes previously linked to neurodevelopmental outcomes: manufactured chemicals, air pollution, and pharmacological, nutrition and sunlight-related factors) and methylation of the *CYP19A1* P1.f promoter using Weighted Quantile Sum (WQS) regression. We found that the WQS mixture index, a weighted combination of the prenatal exposures, was positively associated with higher P1.f methylation (Adjusted Mean Difference (AMD) = 0.712 (95% CI 0.11, 1.315), p = 0.021), indicating reduced brain aromatase activity. Prenatal exposures with the strongest contribution to the mixture effect included bisphenols (including BPA), reduced sunlight, household mould, phthalates, low folate intake, and air pollution. These findings highlight epigenetic modification of the aromatase gene as a biologically plausible, convergent mechanism through which multiple environmental risk factors for autism may exert effects.

## Introduction

Over the past fifty years, the incidence of autism and other child neurodevelopmental disorders has risen sharply.^1^ Autism is a multifactorial condition, with both genetic and environmental causes, and is diagnosed clinically based on restricted, repetitive patterns of behaviour and difficulties in social communication.^2^ The rise in autism is only partly accounted for by changes in diagnostic criteria and improved awareness, and is too rapid to be explained by genetics alone, indicating that modern environmental exposures may be critical.^3^ Notably, several classes of manufactured chemicals identified as prenatal risk factors for autism have increased in prevalence over the same period.^4^

Three observations suggest a central role for hormone-related neurodevelopmental mechanisms in the aetiology of autism. First, autism is diagnosed considerably more often in males than in females.^5^ Second, multiple manufactured chemicals linked to increased autism risk during pregnancy are known endocrine disruptors.^5^ Third, many genes and molecular pathways implicated in autism are involved in endocrine signalling.^6,7^

Key among these is the gene *CYP19A1*, encoding the aromatase enzyme, which converts androgens to estrogens.^5^ Estrogen signalling is essential for early neurodevelopment in both males and females, regulating neuronal growth, connectivity, plasticity, and other key processes.^7^ However, given their substantially lower systemic estrogen and higher testosterone, males are particularly dependent on aromatase to ensure balanced hormonal signalling in the brain. Accordingly, *CYP19A1* is highly expressed in the male brain during foetal development, particularly in the amygdala, a region linked to social and fear-related information processing.^5^ Reduced *CYP19A1* expression has been observed in postmortem brain tissue from individuals with autism when compared to age-matched controls.^8^ Interestingly, these observations align with the “extreme male brain” theory of autism, which posits that autism arises from an imbalance between androgen and estrogen signalling.^7^

Importantly, aromatase activity can be altered by both genetic and environmental factors, including by exposure to endocrine-disrupting chemicals. Epigenetic processes—reversible, and often heritable, changes in gene expression without alteration of the underlying DNA sequence—provide a mechanism by which environmental exposures can influence intrinsic genetic programmes.^9^ Our recent work indicates that prenatal exposure to the plastic chemical bisphenol A (BPA), a well-studied environmental risk factor for autism, can interfere with brain aromatase activity epigenetically by eliciting increased DNA methylation of *CYP19A1* at the brain-specific promoter, P1.f, as measured in cord blood.^5^ Methylation at P1.f shows high concordance between blood and brain (Spearman r = 0.94),^10^ supporting the use of cord-blood measures as a surrogate marker of brain methylation in early development. Across two birth cohorts, higher methylation of the *CYP19A1* P1.f promoter also mediated the impact of prenatal BPA exposure on methylation of the CREB-binding region of the *BDNF* gene, a central regulator of neurodevelopment.^11^ We further reported that both aromatase knockout mice and mice exposed prenatally to BPA show similar neuroanatomical changes, including reduced cortical and amygdalar dendrite length, and autism-like behaviours.^11^

Several additional potential environmental risk factors for autism have been shown to alter aromatase activity, including phthalate plasticizers,^12^ selective serotonin reuptake inhibitors (SSRIs),^13^ and low vitamin D,^14,15^ raising the possibility that aromatase represents one convergent mechanism, particularly in males, linking the prenatal environment to autism symptomology.

As individuals are exposed *in utero* to constellations of environmental factors simultaneously, there is a need for statistical approaches that can capture the cumulative impact of co-occurring exposures on aromatase regulation. Weighted Quantile Sum (WQS) regression is one such approach,^16^ a popular method for estimating the joint effect of correlated exposures while reducing dimensionality and addressing multicollinearity. By assigning weights to each component of the mixture based on their relative contribution to the outcome, WQS regression facilitates both estimation of the overall effect and identification of key drivers within the exposure set.

Here, using data from the Barwon Infant Study (BIS), we use WQS to extend our earlier findings for BPA by investigating the effect of a broader mixture of 25 environmental factors, linked previously to neurodevelopmental outcomes, on methylation of the *CYP19A1* brain promoter P1.f in umbilical cord blood (Figure 1).

**Figure 1.**
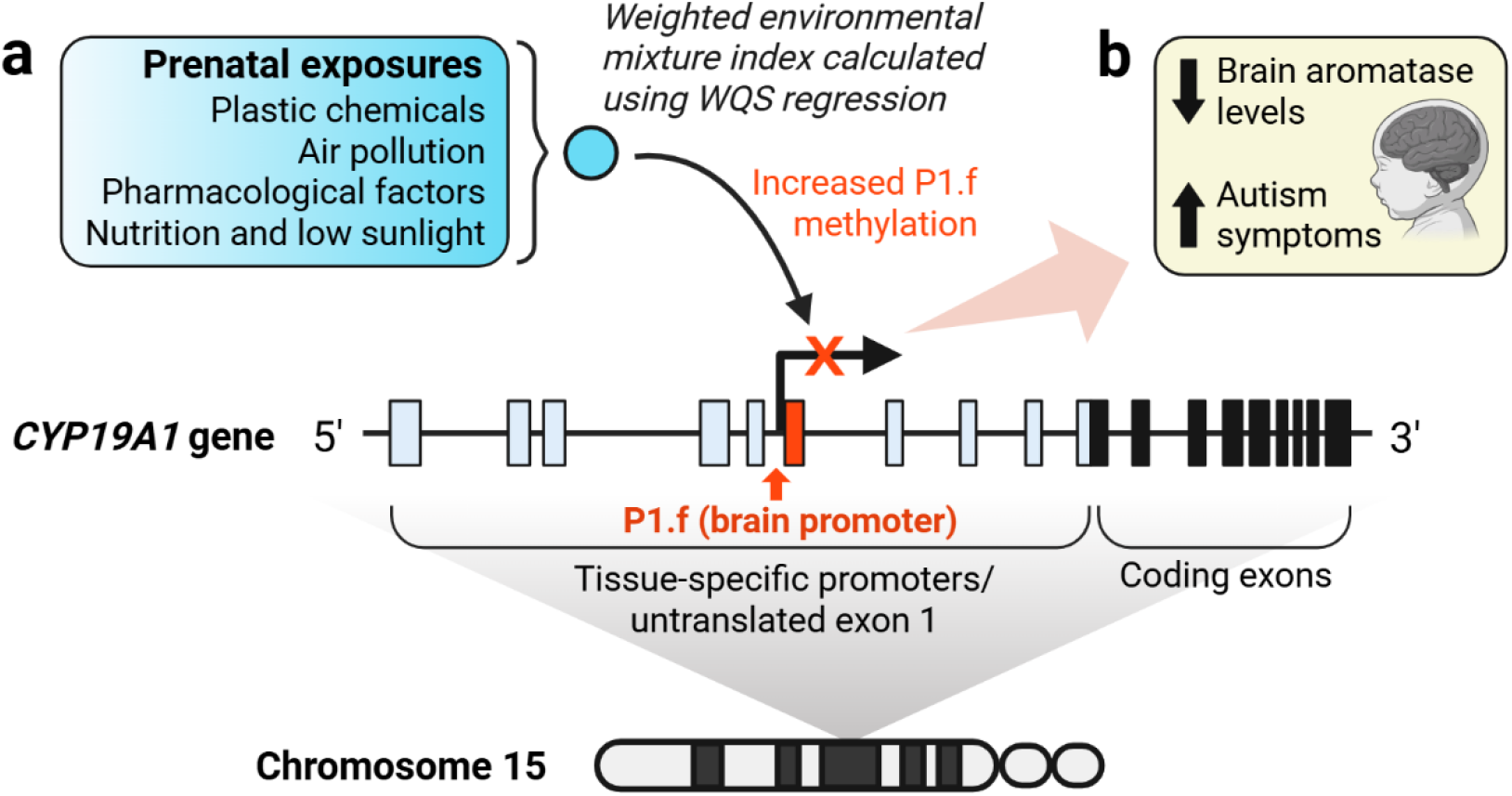
Study overview. **a**. We used Weighted Quantile Sum (WQS) regression to analyse the association between a mixture of 25 environmental exposures, from five classes (shown in the blue box), and DNA methylation of the *CYP19A1* gene, which encodes the aromatase enzyme. *CYP19A1* has a complex regulatory region, with multiple tissue-specific promoters interspersed among the segments of non-coding exon 1.^17^ Activation of a given promoter by transcription factors present in the corresponding tissue type elicits the transcriptional splicing of the adjacent exon-1 segment to the coding exons, resulting in an alternative *CYP19A1* transcript with a unique 5’ end in that tissue type. Here we analysed methylation of the brain-specific promoter, P1.f (depicted by the larger black arrow), which controls *CYP19A1* transcription, and thus aromatase enzyme levels, in the brain. **b**. Previous studies, including our past work, have linked reduced brain aromatase levels to increased likelihood of autism.^10^

## Methods

### Study design and participants

The population-derived BIS birth cohort (mothers, n = 1064; infants, n = 1074; 10 set of twins; n = 906 infants with measures available for this study) was recruited antenatally in the Barwon region of Victoria between June 2010 and June 2013. Exclusion criteria included mothers who were younger than 18 years of age or who required interpreter assistance with questionnaires, and infants who were delivered prior to 32 weeks, who developed serious illness, or who had a genetic disease or major congenital malformation. Further eligibility criteria, population characteristics and measurement details have been previously described.^18^

### Prenatal exposures

**Box 1**. *Summary of all prenatal environmental exposures examined (n = 25)*

**Plastic chemical exposures (n = 11)**

▪ *Low-Molecular Weight Phthalates (LMWP)*: Monomethyl phthalate (MMP), Mono-n-butyl phthalate (MnBP), Mono-isobutyl phthalate (MiBP), and Monoethyl phthalate (MEP)
▪ *High-Molecular Weight Phthalates (HMWP)*: Monobenzyl phthalate (MBzP), Mono-(2-ethyl-5-oxohexyl) phthalate (MEOHP), Mono-(2-ethyl-5-carboxypentyl) phthalate (MECPP), and Mono-(2-ethyl-5-hydroxyhexyl) phthalate (MEHHP)
▪ *Bisphenols*: Bisphenol A (BPA), Bisphenol S (BPS), and Bisphenol F (BPF)

**Outdoor and indoor air pollution (n = 4)**

▪ Two measures of ambient air pollution: Particulate Matter (PM_2.5_) and Nitrogen Dioxide (NO_2_)
▪ Heating (gas/oil/kerosine/diesel) in residence
▪ Mould and/or dampness in residence

**Pharmacological factors (n = 6)**

▪ Selective Serotonin Reuptake Inhibitors (SSRI) or Serotonin and Norepinephrine Reuptake Inhibitors (SNRI) use
▪ Antibiotics use
▪ Paracetamol use (> 6 days per month)
▪ Alcohol use
▪ Two measures of tobacco smoke: maternal smoking and passive smoke exposure

**Nutrition and sunlight (n = 4)**

▪ Folate levels (red blood cell indices)
▪ Fish oil supplementation
▪ UV radiation estimated separately in 1st trimester and 2nd trimester.

Twenty-five prenatal exposures spanning chemical, air pollutant, pharmacological, and nutritional/sunlight-related factors were selected based on previously reported associations with neurodevelopment and data availability in BIS (Box 1).^19-22^ Chemical exposures were measured in maternal urine collected at 36 weeks’ gestation and included eight phthalate metabolites and three bisphenols. Urinary phthalate and bisphenol concentrations were corrected for batch and specific gravity.^23^ For phthalate metabolites, measurements below the limit of detection (LOD) were imputed with 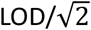.^24^ Phthalate metabolites were additionally corrected for time of day of urine sample collection using the residual method.^25^ Air pollution was estimated from the mother’s postcode at birth. Household mould/dampness at 28 weeks and pharmacological exposures from three months pre-conception to birth were determined from questionnaire data.^19^ Ambient ultraviolet radiation (UVR) exposure was estimated separately for trimesters 1 and 2 from satellite data. Fish oil supplementation was determined from the food frequency questionnaire administered at 28 weeks’ gestation. Red blood cell folate levels were measured at 28–32 weeks’ gestation. See Table 1 caption for further details on timing, and Supplementary Methods for full exposure descriptions and processing procedures.

**Table 1.**
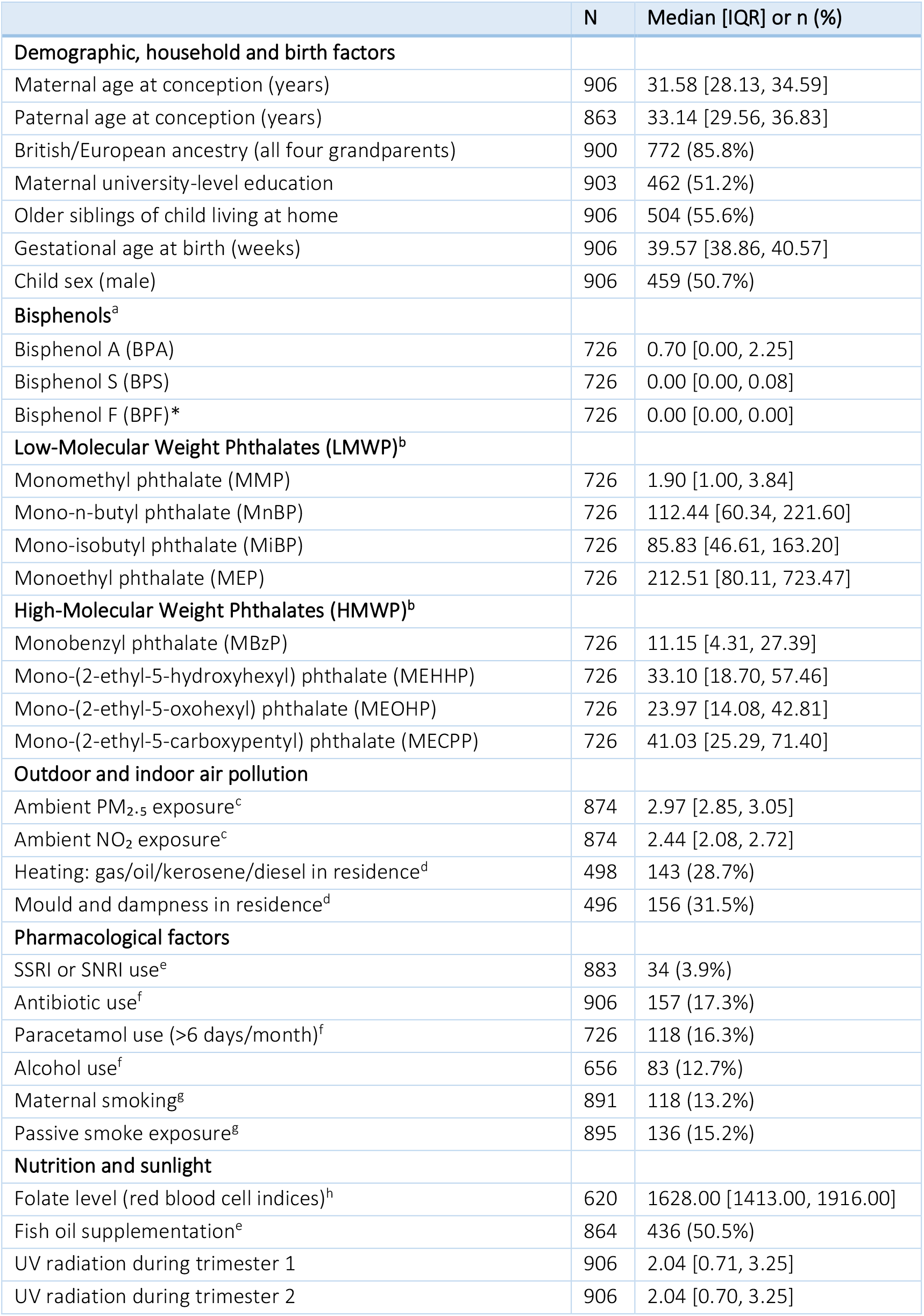
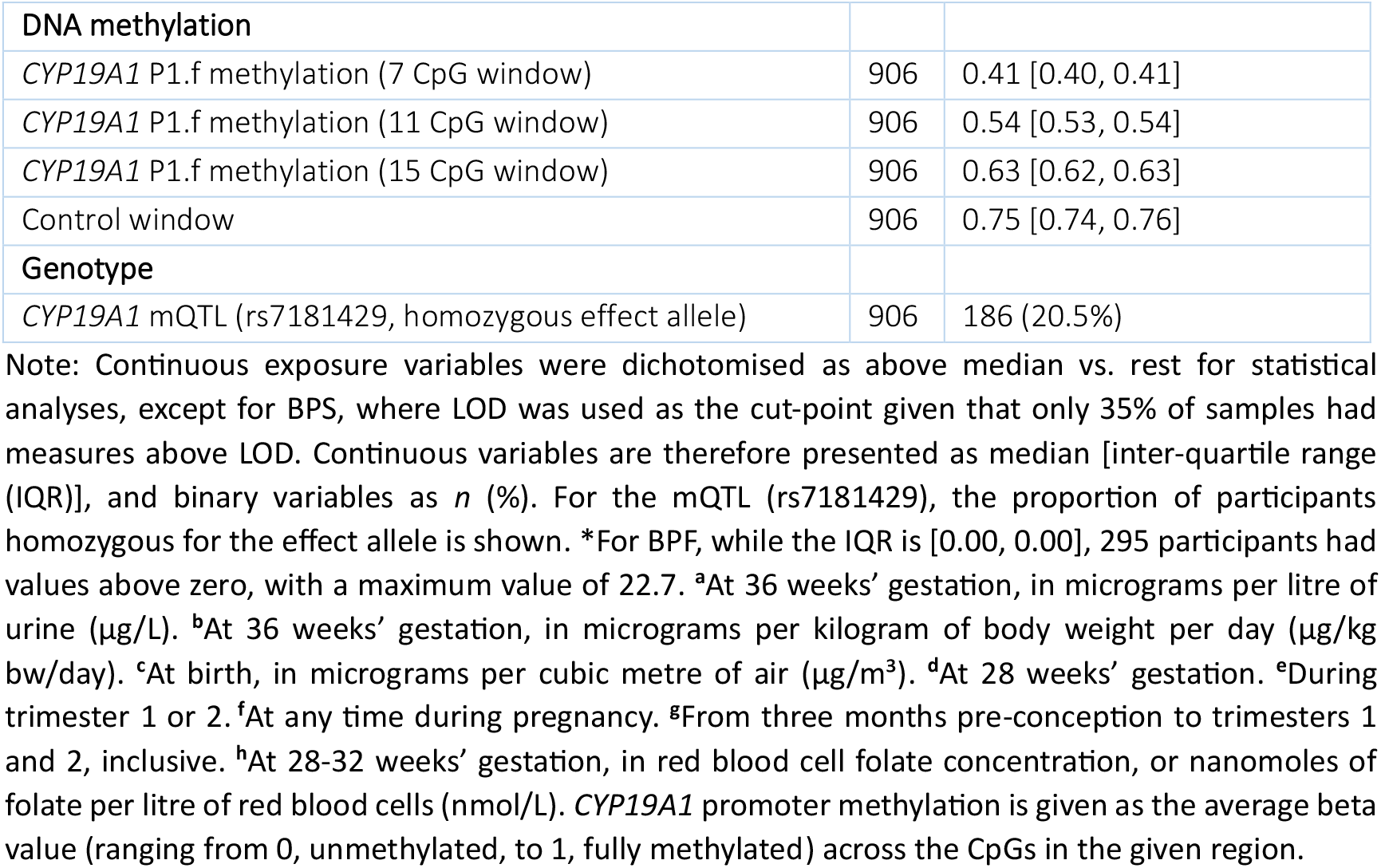
The distribution of key characteristics in the Barwon Infant Study.

|  | N | Median [IQR] or n (%) |
| --- | --- | --- |
| **Demographic, household and birth factors** |  |  |
| Maternal age at conception (years) | 906 | 31.58 [28.13, 34.59] |
| Paternal age at conception (years) | 863 | 33.14 [29.56, 36.83] |
| British/European ancestry (all four grandparents) | 900 | 772 (85.8%) |
| Maternal university-level education | 903 | 462 (51.2%) |
| Older siblings of child living at home | 906 | 504 (55.6%) |
| Gestational age at birth (weeks) | 906 | 39.57 [38.86, 40.57] |
| Child sex (male) | 906 | 459 (50.7%) |
| **Bisphenols<sup>a</sup>** |  |  |
| Bisphenol A (BPA) | 726 | 0.70 [0.00, 2.25] |
| Bisphenol S (BPS) | 726 | 0.00 [0.00, 0.08] |
| Bisphenol F (BPF)* | 726 | 0.00 [0.00, 0.00] |
| **Low-Molecular Weight Phthalates (LMWP)<sup>b</sup>** |  |  |
| Monomethyl phthalate (MMP) | 726 | 1.90 [1.00, 3.84] |
| Mono-n-butyl phthalate (MnBP) | 726 | 112.44 [60.34, 221.60] |
| Mono-isobutyl phthalate (MiBP) | 726 | 85.83 [46.61, 163.20] |
| Monoethyl phthalate (MEP) | 726 | 212.51 [80.11, 723.47] |
| **High-Molecular Weight Phthalates (HMWP)<sup>b</sup>** |  |  |
| Monobenzyl phthalate (MBzP) | 726 | 11.15 [4.31, 27.39] |
| Mono-(2-ethyl-5-hydroxyhexyl) phthalate (MEHHP) | 726 | 33.10 [18.70, 57.46] |
| Mono-(2-ethyl-5-oxohexyl) phthalate (MEOHP) | 726 | 23.97 [14.08, 42.81] |
| Mono-(2-ethyl-5-carboxypentyl) phthalate (MECPP) | 726 | 41.03 [25.29, 71.40] |
| **Outdoor and indoor air pollution** |  |  |
| Ambient PM<sub>2.5</sub> exposure<sup>c</sup> | 874 | 2.97 [2.85, 3.05] |
| Ambient NO<sub>2</sub> exposure<sup>c</sup> | 874 | 2.44 [2.08, 2.72] |
| Heating: gas/oil/kerosene/diesel in residence<sup>d</sup> | 498 | 143 (28.7%) |
| Mould and dampness in residence<sup>d</sup> | 496 | 156 (31.5%) |
| **Pharmacological factors** |  |  |
| SSRI or SNRI use<sup>e</sup> | 883 | 34 (3.9%) |
| Antibiotic use<sup>f</sup> | 906 | 157 (17.3%) |
| Paracetamol use (>6 days/month)<sup>f</sup> | 726 | 118 (16.3%) |
| Alcohol use<sup>f</sup> | 656 | 83 (12.7%) |
| Maternal smoking<sup>g</sup> | 891 | 118 (13.2%) |
| Passive smoke exposure<sup>g</sup> | 895 | 136 (15.2%) |
| **Nutrition and sunlight** |  |  |
| Folate level (red blood cell indices)<sup>h</sup> | 620 | 1628.00 [1413.00, 1916.00] |
| Fish oil supplementation<sup>e</sup> | 864 | 436 (50.5%) |
| UV radiation during trimester 1 | 906 | 2.04 [0.71, 3.25] |
| UV radiation during trimester 2 | 906 | 2.04 [0.70, 3.25] |
| **DNA methylation** |  |  |
| *CYP19A1* P1.f methylation (7 CpG window) | 906 | 0.41 [0.40, 0.41] |
| *CYP19A1* P1.f methylation (11 CpG window) | 906 | 0.54 [0.53, 0.54] |
| *CYP19A1* P1.f methylation (15 CpG window) | 906 | 0.63 [0.62, 0.63] |
| Control window | 906 | 0.75 [0.74, 0.76] |
| **Genotype** |  |  |
| *CYP19A1* mQTL (rs7181429, homozygous effect allele) | 906 | 186 (20.5%) |

Note: Continuous exposure variables were dichotomised as above median vs. rest for statistical analyses, except for BPS, where LOD was used as the cut-point given that only 35% of samples had measures above LOD. Continuous variables are therefore presented as median [inter-quartile range (IQR)], and binary variables as *n* (%). For the mQTL (rs7181429), the proportion of participants homozygous for the effect allele is shown. \*For BPF, while the IQR is [0.00, 0.00], 295 participants had values above zero, with a maximum value of 22.7. <sup>a</sup>At 36 weeks' gestation, in micrograms per litre of urine (µg/L). <sup>b</sup>At 36 weeks' gestation, in micrograms per kilogram of body weight per day (µg/kg bw/day). <sup>c</sup>At birth, in micrograms per cubic metre of air (µg/m<sup>3</sup>). <sup>d</sup>At 28 weeks' gestation. <sup>e</sup>During trimester 1 or 2. <sup>f</sup>At any time during pregnancy. <sup>g</sup>From three months pre-conception to trimesters 1 and 2, inclusive. <sup>h</sup>At 28-32 weeks' gestation, in red blood cell folate concentration, or nanomoles of folate per litre of red blood cells (nmol/L). *CYP19A1* promoter methylation is given as the average beta value (ranging from 0, unmethylated, to 1, fully methylated) across the CpGs in the given region.

### DNA methylation profiling and genotyping

Umbilical cord blood DNA methylation was profiled using the Illumina MethylationEPIC array (over 850,000 CpG sites profiled on each sample).^26^ Quality control, normalization, and probe filtering followed standard procedures as outlined in Supplementary Methods. After pre-processing and filtering, 906 samples and 798,259 probes were retained. Cell-type composition and maternal contamination were estimated as in previous studies^10^ for inclusion as covariates. Aromatase *CYP19A1* brain promoter P1.f maps to flanking positions chr15:51570141-51570603 in human-genome build GRCh37/hg19.^27^ Average methylation across P1.f was assessed principally for a 7 CpG window directly over the promoter, but the specificity of effect was further evaluated using an expanding window approach, beginning with the 7 CpG overlap, then extending upstream to 11 and 15 CpG overlaps. As a control window, average methylation of the entire upstream *CYP19A1* promoter region, excluding the 7 CpG window over P1.f, was used.

Genome-wide genotyping was performed on DNA samples extracted from cord or 12-month whole blood with the Illumina Global Screening Array. Imputation was completed using the Sanger Imputation Server based on the Haplotype Reference Consortium reference panel.^28^ See Supplementary Methods for full details.

### Statistical analysis

To model the joint effect of the mixture of environmental exposures on methylation at the *CYP19A1* brain promoter P1.f, we applied Weighted Quantile Sum (WQS) regression using the *gWQS* R package,^16^ specifying repeated holdout validation (*rh* = 100) and bootstrapping (*b* = 1000) in the call to the gwqs() function. In each of the 100 iterations, data were randomly split 40/60%^29^ into training and validation sets. Variables contributing more than 𝔠 = 0.04 (the reciprocal of the number of exposures, n=25) to the weighted index were considered significant contributors.^30^ WQS regression requires all exposures to be on the same scale. Given that some exposures were categorical (yes/no), we transformed all continuous exposures into binary indicators using the classification of above median (or above LOD) vs. the rest. WQS regression also requires the association between the exposure mixture and the outcome to be constrained in either the positive or the negative direction. Based on prior literature on the association between the 25 risk factors and adverse neurodevelopment,^19-22^ we report results from the positive direction model, evaluating whether higher exposures are associated with increased methylation at the *CYP19A1* P1.f promoter. Results from the negative direction model are provided in the Supplement for comparison. Some participants were missing data on certain environmental exposures and covariates. We imputed missing data using multiple imputation with the *MICE* R package, generating 30 imputed datasets and including in the imputation variables plausibly associated with missingness. We implemented WQS regression on each of the 30 imputed datasets and employed Rubin’s rules^16^ to derive a single pooled estimate for each model. In the WQS and individual regression analyses, adjustment was made for cord-blood blood cell proportions (CD8(+) T cells, CD4(+) T cells, B lymphocyte cells, and granulocytes), maternal contamination of cord blood (yes/no), time of day of urine collection, sex, gestational age, birth year, and a genetic regulator of *CYP19A1* brain promoter methylation (the methylation quantitative trait locus (mQTL) variant rs7181429 associated in the Human Whole Blood mQTL Atlas with methylation of promoter P1f).^29^ See Supplementary Methods for full preprocessing and modelling details.

## Results

### Participant characteristics

Key characteristics of the BIS cohort are presented in Table 1. Maternal exposure to bisphenols and phthalates varied substantially across participants and analytes, with BPA showing a median of 0.70 ng/mL and interquartile (IQR) range of 0.00-2.25 µg/mL, while the low molecular weight phthalate MiBP had a median of 85.83 µg/kg bw/day and IQR of 46.61-163.20 µg/kg bw/day. Air pollution estimates indicated modest levels of ambient PM_2.5_ and NO_2_, with 29% and 32% of participants reporting household use of combustion heating and mould exposure, respectively. Pharmacological exposures were less common; 17% reported antibiotic use and 4% reported SSRI/SNRI use. Approximately half of the participants reported fish oil supplementation, and median UV radiation exposure was comparable across the first and second trimesters. DNA methylation at the *CYP19A1* P1.f promoter exhibited tight distributions, with an average methylation beta value of 0.41 for the 7-CpG window directly over the P1.f promoter. Lastly, 21% of participants were homozygous for the effect allele at the CYP19A1 mQTL (rs7181429).

### Association between individual exposures and aromatase P1.f methylation

Before investigating combined effects, we analysed the association between the 25 exposures individually and methylation of the *CYP19A1* (aromatase) P1.f brain promoter (captured by mean methylation of a 7 CpG window across the promoter) using multiple linear regression (Table 2). Continuous environmental exposures were dichotomised at the median (or LOD) for consistency with the subsequent WQS regression analysis. Adjusting for cord-blood cell-type proportions, maternal contamination of cord blood, time of day of urine collection, sex, gestational age, birth year, and the mQTL genetic variant rs7181429, prenatal BPA exposure was associated with increased *CYP19A1* P1.f methylation (Adjusted Mean Difference (AMD) = 0.15 (95% CI 0.05, 0.25), *p* = 0.004), as reported previously. Prenatal MnBP exposure was also nominally associated with higher P1f methylation (AMD = 0.10 (95% CI 0.001, 0.21), *p* = 0.049). Controlling the false discovery rate at 5% for this investigation of each factor in isolation, BPA alone remained borderline significant (*p* = 0.091). However, 18 of the 25 factors showed a positive direction of effect, indicating that a mixture approach could capture a stronger combined effect.

**Table 2.** Association between individual prenatal environmental exposures and CYP19A1 brain promoter P1.f methylation.

| Prenatal Exposure | Estimate | P-Value | Adjusted P-Value |
| --- | --- | --- | --- |
| High BPA | 0.149 (95% CI 0.049, 0.250) | 0.004 | 0.091 |
| Household mould | 0.124 (95% CI -0.004, 0.251) | 0.060 | 0.360 |
| High MnBP | 0.104 (95% CI 0.001, 0.207) | 0.049 | 0.360 |
| SSRI/SNRI use | 0.101 (95% CI -0.129, 0.331) | 0.380 | 0.702 |
| Antibiotics use | 0.079 (95% CI -0.037, 0.195) | 0.180 | 0.630 |
| High ambient PM<sub>2.5</sub> | 0.076 (95% CI -0.014, 0.167) | 0.090 | 0.432 |
| Paracetamol use | 0.064 (95% CI -0.067, 0.194) | 0.330 | 0.700 |
| Low UVR T1 | 0.057 (95% CI -0.033, 0.147) | 0.210 | 0.630 |
| High MiBP | 0.047 (95% CI -0.053, 0.148) | 0.350 | 0.700 |
| Low UVR T2 | 0.035 (95% CI -0.053, 0.123) | 0.430 | 0.737 |
| Low folate | 0.030 (95% CI -0.074, 0.135) | 0.560 | 0.747 |
| No fish oil | 0.027 (95% CI -0.064, 0.118) | 0.550 | 0.747 |
| High ambient NO<sub>2</sub> | 0.015 (95% CI -0.075, 0.104) | 0.740 | 0.910 |
| Fuel-based heating | 0.011 (95% CI -0.125, 0.147) | 0.870 | 0.910 |
| High MEOHP | 0.009 (95% CI -0.088, 0.106) | 0.850 | 0.910 |
| High MEP | 0.005 (95% CI -0.095, 0.106) | 0.910 | 0.910 |
| High MBzP | 0.005 (95% CI -0.094, 0.104) | 0.910 | 0.910 |
| High MECPP | -0.012 (95% CI -0.109, 0.084) | 0.800 | 0.910 |
| High MEHHP | -0.034 (95% CI -0.130, 0.062) | 0.480 | 0.747 |
| High BPF | -0.045 (95% CI -0.187, 0.096) | 0.520 | 0.747 |
| High MMP | -0.052 (95% CI -0.148, 0.044) | 0.290 | 0.696 |
| Active maternal smoking | -0.069 (95% CI -0.192, 0.055) | 0.270 | 0.696 |
| Passive maternal smoke exposure | -0.085 (95% CI -0.217, 0.048) | 0.200 | 0.630 |
| Alcohol Use | -0.191 (95% CI -0.351, -0.032) | 0.019 | 0.228 |

Each association was tested using linear regression, with adjustment for cord-blood blood cell-type proportions, maternal contamination of cord blood, time of day of urine collection, sex, gestational age, birth year, and the mQTL genetic variant rs7181429. Continuous exposures were dichotomised at the median (or LOD), with values above these thresholds designated as “high”. P-values in the rightmost column are adjusted for multiple comparisons using the Benjamini-Hochberg procedure.

### Association between prenatal environmental mixture and aromatase P1.f methylation

Accordingly, we employed WQS regression to estimate the aggregate effect of the mixture of 25 prenatal exposures on *CYP19A1* P1.f methylation and obtain a ranking of exposures by their contribution to this effect (Figure 2). Adjusting for the same covariate set (see Table 2 footnote), the resulting environmental mixture score (which can range from 0 to 1) was associated with increased P1.f methylation (AMD_WQS_ = 0.71 (95% CI 0.11, 1.32), *p* = 0.021) (Figure 2a). The AMD_WQS_ of 0.71 reflects the estimated increase in *CYP19A1* P1.f methylation (in averaged beta values) associated with a full-unit increase in the WQS index—that is, moving from having no high exposures to a profile in which all exposures are in the high (above median or LOD) category. In practice, this represents the effect of increasing cumulative high-exposure burden across the weighted mixture, with greater influence from the most strongly weighted exposures. Exposures contributing most strongly to the mixture index were low ultraviolet light exposure in trimester 2, BPA exposure, and exposure to household damp mould, with relative weights of 0.128, 0.128 and 0.097 respectively (Figure 2b). In total, nine of the 25 exposures were assigned weights above the threshold of 𝔠 = 0.04 reflecting a material contribution to the association with P1.f methylation, with the additional six exposures being BPF, the phthalate plasticizer MiBP, low folate, two indicators of increased air pollution (PM_2.5_ and NO_2_) and low ultraviolet light exposure in trimester 1.

**Figure 2.**
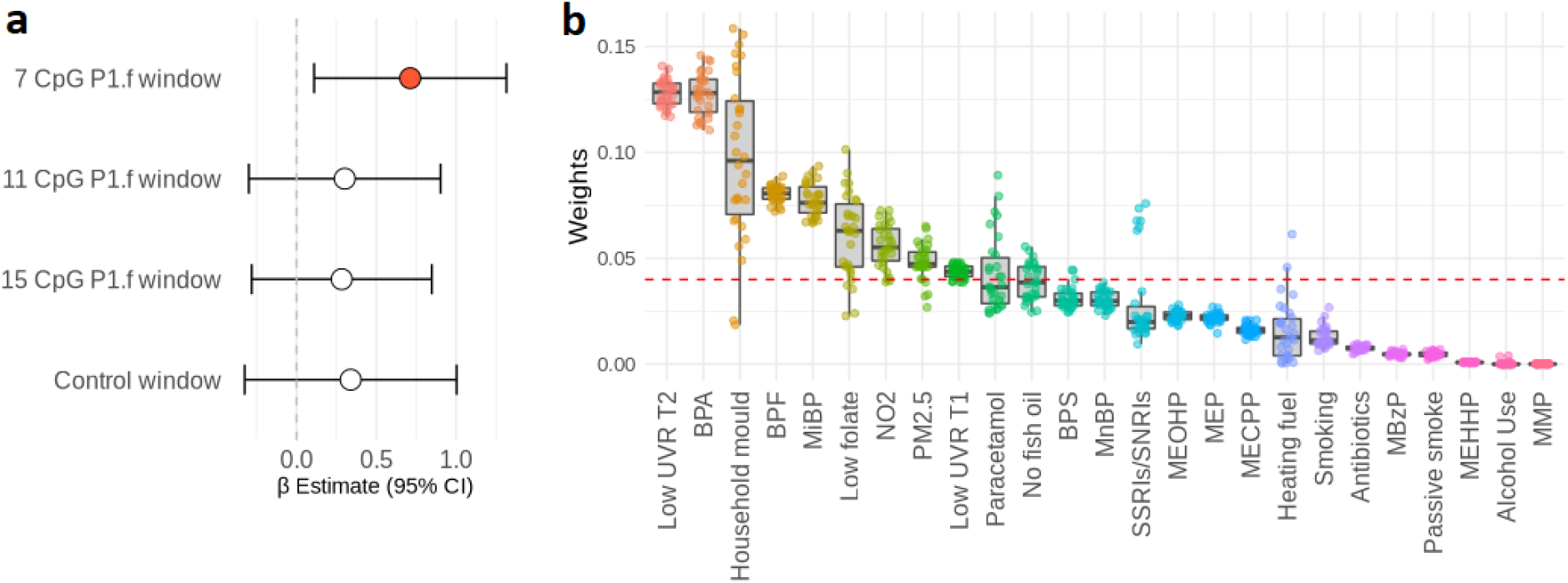
Analysis of the association between a mixture of 25 prenatal environmental factors and CYP19A1 (Aromatase) brain promoter P1.f methylation using Weighted Quantile Sum (WQS) regression. **a**. Association between the WQS mixture score and methylation of the 7 CpG window directly over the brain-specific P1.f promoter, 11 and 15 CpG windows including P1.f but extended upstream, and a control window upstream of, but not including, P1.f. Adjustment was made for cord-blood blood cell-type proportions, maternal contamination of cord blood, time of day of urine collection, sex, gestational age, birth year, and the mQTL genetic variant rs7181429. These findings indicate a combined effect of the prenatal environmental factors specifically at the *CYP19A1* brain promoter P1.f. **b**. Weights assigned to each environmental exposure in the WQS model for the 7-CpG P1.f window across 30 imputations, reflecting their individual contributions to the mixture score. Each point represents the weight assigned to the exposure by the WQS model in a specific imputed dataset, with the median pooled weight across all imputations given by the centre bar in each boxplot. Exposures assigned weights above the dotted red line (𝔠 = 0.04) showed a significant contribution to the mixture effect.

To evaluate the specificity of this effect on the P1.f brain promoter, we next used WQS regression to test the association between the mixture of 25 chemicals and mean methylation of (i) an 11 CpG window and (ii) a 15 CpG window, both upstream extensions of the 7 CpG P1.f window, as well as (iii) a control window encompassing the entire *CYP19A1* promoter region upstream of, but not including, P1.f (Figure 2a). The resulting WQS mixture scores were not associated with methylation of these regions (11 CpG window: AMD_WQS_ = 0.302 (95% CI -0.299, 0.902), *p* = 0.324; 15 CpG window: AMD_WQS_ = 0.283 (95% CI -0.281, 0.847), *p* = 0.325; control window: AMD_WQS_ = 0.339 (95% CI -0.325, 1.003), *p* =0.316). Moreover, corresponding negative direction WQS models—which test whether exposure to the mixture is associated with decreased methylation at each *CYP19A1* methylation window—were nonsignificant, reinforcing the primary positive direction results (Supplementary Table 2).

These findings support the hypothesis that the aromatase gene is a convergent hub through which multiple neurodevelopmentally relevant environmental factors operate in concert with BPA exposure and that these effects are specific to the P1.f brain promoter.

## Discussion

In an extensively profiled population-based birth cohort, we found evidence that multiple environmental exposures act together to increase methylation of the *CYP19A1* brain promoter, P1.f. This promoter regulates brain levels of the aromatase enzyme, a critical determinant of neurodevelopment. Disruption of the aromatase enzyme is a key mechanism through which prenatal exposure to the plastic chemical BPA increases autism risk in males, as we have previously shown.^5^

Here, we investigated an environmental mixture consisting of 25 factors previously linked to neurodevelopmental outcomes, including autism.^19-22^ The top five factors contributing to *CYP19A1* P1.f methylation included two bisphenols (BPA and BPF) and a phthalate metabolite (MiBP), demonstrating that plastic chemicals remain central to aromatase programming in this broader mixture effect. The MiBP finding is consistent with previous studies implicating di-(2-ethylhexyl) phthalate (DEHP) and its metabolite mono-2-ethylhexyl phthalate (MEHP) in aromatase suppression in neuronal cell culture.^31-33^ Strong contributors also included low prenatal UVR in trimester 2, higher household mould exposure, low folate intake, and higher air pollution exposure.^34^ We have previously linked low gestational UVR to offspring neuroinflammatory disease.^35^ Rodent studies have already demonstrated that developmental vitamin D deficiency *in utero* increases methylation of the *CYP19A1* P1f promoter. ^36^ UVR exposure may be a proxy for low prenatal vitamin D levels in this setting.^37^ While prenatal SSRI usage did not meet the threshold of 𝔠 = 0.04 in the pooled WQS weights, the high weights assigned in a subset of imputations are consistent with *in vitro* assays and other studies linking multiple SSRI drugs to decreased aromatase enzyme levels.^38,39^

There was no association between the mixture of 25 environmental factors and average methylation of larger 11- and 15-CpG methylation windows extended upstream from the brain promoter or of the entire upstream promoter excluding P1.f. Instead, the finding was specific to the 7-CpG methylation window directly over the brain promoter, highlighting a targeted effect on brain aromatase activity. While P1.f methylation was measured in cord blood, human whole blood and brain aromatase methylation patterns are highly correlated.^5^ The association with hypermethylation of the brain promoter persisted after adjustment for an mQTL (genetic variant) linked to methylation of the 7 CpG window and for genetic ancestry. Genetic confounding is therefore unlikely to explain this association. We also adjusted for any maternal contamination of cord blood.

Several mechanisms may account for these convergent effects on the *CYP19A1* brain promoter. First, the majority of the 25 factors are known to elevate reactive oxygen species (ROS),^20,40,41^ shifting the natural cellular balance between pro- and antioxidant activity towards oxidative stress. In particular, hydrogen peroxide (H_2_O_2_), a key ROS, has been shown in animal models to impair estrogen signalling, leading specifically to reduced brain aromatase expression.^42^ Second, many of the exposures— particularly the plastic chemicals among the top contributors—are endocrine disruptors that interfere directly with the estrogen receptors ERa and ERb.^43^ Notably, ERa regulates *CYP19A1* (or cyp19a1 in mouse cell lines) specifically via the brain promoter P1.f, with little evidence of activity at *CYP19A1’s* other promoters.^11^ Both ERa and ERb recruit epigenetic modifiers that alter methylation of target promoters upon binding. Thus, inhibition of these receptors by bisphenols and other endocrine disruptors would reduce P1.f binding and increase its methylation, consistent with our findings.^44^ Together, these unifying mechanisms plausibly explain both the mixture effect on *CYP19A1* methylation as well as its specificity for the brain promoter, P1.f. Aromatase down-regulation subsequently elicits changes across key neurodevelopmental signalling networks, including those controlled by brain derived neurotrophic factor (*BDNF*) and RAR-related orphan receptor alpha (*RORA*).^10^

Previous work by ourselves^5^ and others^8,45-48^ indicates that, while there are no sex differences in the association between BPA exposure and *CYP19A1* methylation, the consequences of aromatase suppression are more detrimental in males. Male-specific changes identified in aromatase-knockout mouse models include reduced *BDNF* expression in the amygdala,^5,49^ altered cortical EEG activity, neuroanatomical changes (dendrite stunting and sparser spine density), and impaired sociability.^5^ Thus, it appears that the aromatase gene is similarly vulnerable to epigenetic suppression across sexes but the consequences are more adverse for males.^10^

A key strength of this study is the use of WQS regression to investigate the effect of multiple environmental exposures on *CYP19A1* brain promoter methylation. WQS provides a single interpretable index capturing the overall mixture effect, with higher weights assigned to exposures more strongly associated, here, with brain promoter methylation. Mixture approaches are better aligned with real-world exposures, where individuals are not exposed to single pollutants in isolation but to a multiplicity of factors simultaneously.^50^ Mixture methods can detect subtle coordinated effects that may not be evident at the level of single exposures. Moreover, while tolerable daily intake has historically been evaluated for individual chemicals in isolation, public health regulatory bodies are increasingly moving to re-assess safe levels based on real-world mixture effects.^51^ Further strengths include extensive prenatal assessments in the BIS cohort, which allowed us to consider nutritional deficiencies and pharmacological agents that may be important in addition to chemicals and other pollutants. Limitations include the necessarily observational nature of human studies on harmful pollutants. We aimed to address this by careful adjustment for potential confounding factors, but results should be interpreted with some caution. Further, while the WQS framework is ideally suited to examining large numbers of heterogeneous exposures, it does not accommodate non-linear effects or interactions between exposures. Future studies could explore the use of Bayesian Kernel Machine Regression (BKMR) or similar approaches on smaller subsets of exposures.

In summary, these findings build on our recent multimodal study on BPA, aromatase and autism by providing evidence that multiple prenatal exposures—not only BPA—elicit epigenetic changes at the *CYP19A1* brain promoter P1.f. While previous work has largely examined individual exposures in isolation, our use of mixture modelling reveals the cumulative nature of real-world exposures, with other modifiable exposures, including low UVR and increased household mould, phthalate exposure and air pollution, contributing to P1.f methylation alongside BPA. Given the importance of aromatase in male neurodevelopment and its dysregulation in autism and other neurodevelopmental disorders,^52^ these findings underscore the need to re-evaluate environmental safety thresholds for relevant chemicals through the lens of mixture effects. By identifying specific modifiable prenatal chemical, nutritional, and lifestyle factors of concern, this study suggests an actionable approach to addressing the growing burden of neurodevelopmental disorders.

## Author Contributions

S.Ta.: Conceptualisation, Methodology, Data Curation, Formal Analysis, Writing – original draft, Writing –review and editing; K.D.: Formal Analysis, Data Curation, Writing – original draft, Writing – review and editing; S.Th.: Methodology, Data Curation, Writing – review and editing; K.V.: Conceptualisation, Writing – review and editing; C.S.: Conceptualisation, Funding Acquisition, Writing – review and editing; B.N.: Methodology, Data Curation, Writing – review and editing; T.M.: Methodology, Data Curation, Writing – review and editing; M.O.H.: Methodology, Writing – review and editing; R.S., M.T., P.S., P.V.: Conceptualisation, Funding Acquisition, Writing – review and editing; W.C.B.: Conceptualisation, Methodology, Writing – review and editing; D.P. and C.H.Y.: Methodology, Writing –review and editing; A.-L.P.: Conceptualisation, Methodology, Funding Acquisition, Writing – original draft, Writing – review and editing; All Authors: Writing – review and editing.

## Acknowledgements

The authors thank the BIS participants for the generous contribution they have made to this project. The authors also thank current and past staff for their efforts in recruiting and maintaining the cohort and in obtaining and processing the data and biospecimens. We acknowledge Barwon Health, Murdoch Children’s Research Institute, and Deakin University for their support in the development of this research. The other members of the BIS Investigator Group are David Burgner, Lawrence Gray, Len Harrison, and Sarath Ranganathan. We thank John Carlin, Amy Loughman, Fiona Collier, Terry Dwyer, and Katie Allen for their past work as BIS investigators. We thank Andrea Gogos, Billi Newton, and Alicia Bjorksten for manuscript preparation.

