## Supplement for "Prenatal Environmental Determinants of Aromatase Brain-Promoter Methylation in Cord Blood: Chemical, Airborne, Pharmacological, and Nutritional Factors"

[1] The Florey Institute of Neuroscience and Mental Health, The University of Melbourne, Parkville, VIC, Australia
[2] Melbourne Bioinformatics, The University of Melbourne, Parkville, VIC, Australia
[3] Murdoch Children's Research Institute, Royal Children's Hospital, and Department of Paediatrics, The University of Melbourne, Parkville, VIC, Australia
[4] Deakin University, IMPACT – the Institute for Mental and Physical Health and Clinical Translation, School of Medicine, Geelong, VIC, Australia
[5] Faculty of Health, Deakin University, Geelong, VIC, Australia
[6] Children's Health and Environment Program, Child Health Research Centre, The University of Queensland, South Brisbane, QLD, Australia
[7] Barwon Health, Geelong, VIC, Australia
[8] Department of Biochemistry and Pharmacology, The University of Melbourne, Parkville, VIC, Australia
[9] Minderoo Foundation, Perth, WA, Australia

### Contents

**Supplementary methods**

Prenatal exposures

SNP genotyping

DNA methylation profiling

**Supplementary results**

Supplementary Table 1. Positive direction Weighted Quantile Sum (WQS) regression model.

Supplementary Table 2. Negative direction Weighted Quantile Sum (WQS) regression model.

### Supplementary methods

#### Prenatal exposures

**Chemical exposures**: A single spot urine specimen was collected at 36 weeks of gestation, divided into aliquots and frozen at −80 °C before being shipped on dry ice to the Queensland Alliance for Environmental Health Science (QAEHS). There, samples were analysed for 14 phthalate metabolites and five bisphenol analogues using high-performance liquid chromatography/tandem mass spectroscopy with direct injection. QAEHS procedures for the measurements of the phthalate metabolites have been published previously and for the bisphenols.[1,2] After excluding phthalate metabolites with >30% of measurements across the cohort below the limit of detection (LOD), eight phthalate metabolites were used in this study: monoethyl phthalate (MEP), monomethyl phthalate (MMP), mono-n-butyl phthalate (MnBP), monoisobutyl phthalate (MiBP), monobenzyl phthalate (MBzP), mono-(2-ethyl-5-hydroxyhexyl) phthalate (MEHHP), mono-(2-ethyl-5-oxohexyl) phthalate (MEOHP) and mono(5-carboxy-2-ethylpentyl) phthalate (MECPP). Bisphenols were excluded if >90% of measurements across the cohort were below the LOD. Bisphenol-A (BPA), bisphenol-S (BPS) and bisphenol-F (BPF) were used in this study.

**Outdoor and indoor air pollution**: Outdoor ambient pollution in pregnancy in BIS has been previously published.[3] In brief, residential address was used to estimate satellite-based nitrogen dioxide and PM2.5 at each infant’s residential address at birth (2010–2013). Household information including mould, dampness, and heating types for the current and previous residence was collected by questionnaire during the in-clinic study visit at two years. Data relevant to the household the mother occupied pregnancy was only used in this analysis.

**Pharmacological factors**: For active tobacco smoke exposure, we consider any maternal tobacco smoking (yes vs. no) across the 3-month pre-conception period, and the 1$^{st}$ and 2nd trimesters. Information on antidepressant use, including selective serotonin reuptake inhibitors (SSRIs) and serotonin and norepinephrine reuptake inhibitors (SNRIs) was collected in the medication questionnaire during the 1st and 2nd trimester. Paracetamol use (6 + days per month), alcohol consumption, and antibiotic usage was documented for any time during pregnancy.

**Nutrition and sunlight**: UVR dose exposure variables were modelled on previous work by Molloy *et al*.[4] Questionnaire data quantifying exposure to direct sunlight daily were recorded during trimesters 1 and 2 of pregnancy. Ambient UVR was estimated using monthly averages of daily total ambient UVR in standard erythemal doses for Melbourne, latitude 37.5S, using data provided by the Australian Radiation Protection and Nuclear Safety Agency. UVR dose in pregnancy and the first postnatal year was then estimated by multiplying exposure to direct sunlight in hours and regional monthly averages of daily ambient UVR exposure for the time of enquiry. Fish oil supplementation was determined from the food frequency questionnaire administered at 28 weeks’ gestation. Red blood cell folate levels were measured at 28-32 weeks’ gestation.

#### SNP genotyping

DNA samples were extracted from cord and 12-month whole blood using the QIAamp kit (QIAGEN, Hilden, Germany) according to manufacturer's instructions and stored at -80° C. Whole-genome genotyping was performed with Illumina Global Screening Array (Illumina, San Diego, CA, USA). Imputation was completed using the Sanger Imputation Server (Wellcome Sanger Institute, Hinxton, UK) based on the Haplotype Reference Consortium reference panel.[5] Infants were excluded for quality control if initial genotyping was unsuccessful at more than 5% of SNPs. SNPs were dropped if (i) genotyping failed across more than 5% of infants, (ii) minor allele frequency was less than 0.01 or differed by more than 0.2 from the reference population, or (iii) the SNPs were not in Hardy–Weinberg equilibrium.[6]

#### DNA methylation profiling

The Illumina Infinium MethylationEPIC BeadChip (referred to from now as 'EPIC array') was used for DNA methylation profiling of umbilical cord blood from the BIS cohort. Genomic DNA (200 to 500 ng) from cord blood was randomized into 96-well plates and sent to the Kobor laboratory (Canada) for sodium bisulfite treatment and processing on the EPIC array. The EPIC array measures DNA methylation level at more than 850,000 CpG sites (referred to as 'EPIC probes'), and covers all gene promoters, gene bodies and ENCODE-assigned distal regulatory elements. Raw IDAT files were processed and analyzed using the MissMethyl and minfi packages for R, both available from Bioconductor. Samples were checked for quality and those with a mean detection p-value of >0.01 were removed (128 samples), leaving 946 cord blood samples for analysis. Data were normalized for both within and between array technical variation using SWAN (Subset quantile Within Array Normalization). Probes with poor average quality scores (detection *P*-value > 0.01) and cross-reactive probes were removed from further analysis. This left a total of 798,259 probes for cord blood analysis. Cell composition was determined using the estimateCellCounts tool, with the 'CordBlood' reference data used for neonatal blood spot analysis. Maternal contamination of infant cord blood was assessed using a CpG signature.[7]

### Supplementary results

**Supplementary Table 1.** *Positive direction Weighted Quantile Sum (WQS) regression model.*

| ***CYP19A1 Region*** | **Pooled Mean** | **CI Lower** | **CI Upper** | **P-value** |
|---|---|---|---|---|
| *7 CpG P1.f window* | ***0.712*** | ***0.110*** | ***1.315*** | ***0.021**** |
| *11 CpG P1.f window* | 0.302 | -0.299 | 0.902 | 0.324 |
| *15 CpG P1.f window* | 0.283 | -0.281 | 0.847 | 0.325 |
| *Control promoter window* | 0.339 | -0.325 | 1.003 | 0.316 |

Test of a **positive** association between the mixture of 25 prenatal environmental factors and *CYP19A1* (aromatase) brain promoter P1.f DNA methylation (i.e. test for **increased** methylation at this in the presence of the exposures, implying reduced brain aromatase activity). The association was evaluated for average methylation across a 7 CpG window directly over the brain-specific P1.f promoter, 11 and 15 CpG windows including P1.f but extended upstream, and a control window upstream of, but not including, P1.f. Adjustment was made for cord-blood blood cell-type proportions, maternal contamination of cord blood, time of day of urine collection, sex, gestational age, birth year, and the mQTL genetic variant rs7181429.

**Supplementary Table 2.** *Negative direction Weighted Quantile Sum (WQS) regression model.*

| ***CYP19A1 Region*** | **Pooled Mean** | **CI Lower** | **CI Upper** | **P-value** |
|---|---|---|---|---|
| *7 CpG P1.f window* | 0.077 | -0.532 | 0.685 | 0.804 |
| *11 CpG P1.f window* | -0.084 | -0.630 | 0.463 | 0.764 |
| *15 CpG P1.f window* | -0.010 | -0.533 | 0.513 | 0.971 |
| *Control promoter window* | -0.230 | -0.818 | 0.358 | 0.442 |

Test of a **negative** association between the mixture of 25 prenatal environmental factors and *CYP19A1* (aromatase) brain promoter P1.f DNA methylation (i.e. test for **reduced** methylation at this site in the presence of the exposures, implying increased brain aromatase activity). The association was evaluated for average methylation across a 7 CpG window directly over the brain-specific P1.f promoter, 11 and 15 CpG windows including P1.f but extended upstream, and a control window upstream of, but not including, P1.f. Adjustment was made for cord-blood blood cell-type proportions, maternal contamination of cord blood, time of day of urine collection, sex, gestational age, birth year, and the mQTL genetic variant rs7181429. No negative-direction models reached significance.
